# Video-based automated analysis of MDS-UPDRS III parameters in Parkinson disease

**DOI:** 10.1101/2022.05.23.493047

**Authors:** Gaëtan Vignoud, Clément Desjardins, Quentin Salardaine, Marie Mongin, Béatrice Garcin, Laurent Venance, Bertrand Degos

**Author notes:** **Corresponding authors:** Bertrand Degos, Laurent Venance. **Data availability:** The data are not available for public access because of patient privacy concerns but are available from the corresponding author BD on reasonable request.

## Abstract

**Background:** Among motor symptoms of Parkinson’s disease (PD), including rigidity and resting tremor, bradykinesia is a mandatory feature to define the parkinsonian syndrome. MDS-UPDRS III is the worldwide reference scale to evaluate the parkinsonian motor impairment, especially bradykinesia. However, MDS-UPDRS III constitutes an agent-based score making reproducible measurements and follow-up challenging.

**Objectives:** Using a deep learning approach, we developed a tool to compute an objective score of bradykinesia based on the gold-standard MDS-UPDRS III.

**Methods:** In the Movement Disorder unit of Avicenne University Hospital, we acquired a large database of videos of parkinsonian patients performing MDS-UPDRS III protocols. We applied two deep learning algorithms to detect a two-dimensional (2D) skeleton of the hand composed of 21 predefined points, and transposed it into a three-dimensional (3D) skeleton.

**Results:** We developed a 2D and 3D automated analysis tool to study the evolution of several key parameters during the protocol repetitions of the MDS-UPDRS III. Scores from 2D automated analysis showed a significant correlation with gold-standard ratings of MDS-UPDRS III, measured with coefficients of determination for the tapping (0.609) and hand movements (0.701) protocols using decision tree algorithms. The individual correlations of the different parameters measured with MDS-UPDRS III scores carry meaningful information and are consistent with MDS-UPDRS III guidelines.

**Conclusion:** We developed a deep learning-based tool to reliably score and analyze bradykinesia for parkinsonian patients.

## INTRODUCTION

The diagnosis of Parkinson’s disease (PD) is based on the presence of a parkinsonian syndrome, i.e. the association of rigidity and/or rest tremor to bradykinesia, this latter being mandatory for the diagnosis [1]. Bradykinesia is defined as a motor slowness associated with a decrease in the amplitude and/or speed of movement [2–4]. Currently, the evaluation of motor impairment of PD is based on part III (motor) of the Unified Parkinson’s Disease Rating Scale (MDS-UPDRS) composed of 18 items rated from 0 (normal) to 4 (severe), which assess the severity of bradykinesia, rigidity and tremor [5,6]. This score is worldwide-used for patient follow-up in outpatient clinic but also in clinical research and more specifically in therapeutic trials. However, the semi-quantitative assessment of parkinsonian symptoms by the MDS-UPDRS III score suffers from a certain subjectivity and the reproducibility of the measurements arising from assessors is questionable especially in case of non-parkinsonian expertise [7–13]. This may contribute to the difficulty in the follow-up of PD patients and also to induce some biases in clinical research due to the multiplicity of assessors and clinical centers, and variability across longitudinal iterative visits.

Although digital tools have been recently developed in an attempt to improve bradykinesia scoring by providing quantitative measures [14–16], they exhibit limited use in practice because of wearable sensors on patients [17] in a dedicated room or in specific conditions [15,18,19]. More recent studies aimed to circumvent these material issues by developing video-based only tools. First, Park et al. investigated the utility of machine learning-based automatic rating for rest tremor and finger tapping to measure bradykinesia [20]. They showed that it was feasible, reliable when compared to movement disorders specialist ratings and more accurate than a non-trained one. The authors based their analysis on the periodicity of the repeating task videotaped, i.e. the frequency of the movement, to predict a severity score, but the outcomes did not fully reflect the MDS-UPDRS score in which decreases in speed and amplitude of movement were also considered [20]. Another work assessed the global PD severity applying deep learning approach to automatically score the severity level by compiling 7 of the 18 items of the MDS-UPDRS III [21]. Finally, Monje et al. investigated MDS-UPDRS III bradykinesia upper limbs tasks but with the constraint of fixed webcam recordings and with different assessment guidelines from MDS-UPDRS[22]. Although these studies bring interesting approaches, it remains to develop tools to closely establish medically-relevant metrics for a better assessment of bradykinesia.

In this emerging context of computer-based tools and given the importance of bradykinesia evaluation in the PD diagnosis, we developed a tool based on hand-pose estimation and movement analysis to compute an objective score of bradykinesia with multiple medically-relevant parameters using a deep learning approach in accordance with MDS-UPDRS III recommendations [23].

## MATERIAL AND METHODS

### Participants and videos

We included only PD patients according to MDS clinical criteria [1], with the exception of one MSA patient [24] and another one with undetermined atypical parkinsonian syndrome. All patients were consecutively seen in an outpatient clinic by a movement disorder specialist (BD) at Avicenne University Hospital between June 2019 and December 2020. All patients were video-recorded, using standard smartphones (rescaled to 720×1280 pixels with 30 or 60 fps), for the items 3.4, 3.5 and 3.6 of the MDS-UPDRS III corresponding to the assessment of finger tapping, hand movements, and pronation-supination movements of the hands, respectively.

The videos were then stored in a secure database, according to the French data protection authority (Commission Nationale de l’Informatique et des Libertés) recommendations. All participants gave their written informed consents for videos and their analyses. The study was approved by the local ethics committee (CLEA-2019-83) and registered in ClinicalTrials.gov (NCT04974034).

### MDS-UPDRS III ratings of the video recordings

Two movement disorders specialists certified for MDS-UPDRS (BD, MM) and three neurologists trained for movement disorders, but not certified for MDS-UPDRS scoring (CD, QS and BG), rated all the videos for both hands according to the items 3.4, 3.5 and 3.6 of the MDS-UPDRS III [5].

### Hand-pose estimation using deep learning algorithms

We analyzed videos using two deep learning algorithms in order to extract interesting temporal features of the movement. We used a first network, DeepLabCut [23], to extract predefined points of interest in 2D from images. Using an associated software (http://www.mackenziemathislab.org/deeplabcut), we roughly labelled 5 images per video with 21 different points (5 for each finger, and one for the wrist) and the network was trained to detect the 21 joints [25] using the DeepLabCut toolbox. After the 2D hand coordinates detection, several algorithms (see next section) were applied to filter and smooth the movement trajectories. We defined a bounding box around the patient’s hand from the 2D coordinates obtained with DeepLabCut for each timestep, and each frame was cropped along this box. The cropped image was processed through a second network, HandGraphCNN [26], which without specific training predicted 2D and 3D positions of the 21 points. We then processed and analyzed all trajectories (2D DeepLabCut, 2D & 3D HandGraphCNN) to study the temporal evolution of the movement for each protocol. We realized training and inference using Python 3.X, tensorflow and Pytorch on a GPU Nvidia Geforce GTX 1080 Ti.

### Processing of 2D and 3D coordinates

The 2D trajectories extracted from DeepLabCut went through different post-processing processes: (i) using an iterative algorithm, outlier points, where the speed of movement was above a defined threshold, were deleted; (ii) using the score maps for each point given by DeepLabCut, we took out all coordinates for which the probability was below a defined threshold (fixed at p=0.8); (iii) the trace was smoothed using a Savitzky-Golay filter (with parameters window_length=11, polyorder=5) from the *scipy* library. For (i) and (ii), we used linear interpolation to infer the coordinates of deleted points, with *numpy interp* function.

For the 2D and 3D coordinates from HandGraphCNN, we reproduced step (i) and (iii) of the post-processing algorithm.

**Analysis of Bradykinesia protocols using 2D and 3D coordinates**

Using the 2D and 3D trajectories, we computed different metrics, specific for each protocol in order to evaluate bradykinesia symptoms.

- Finger tapping (using 2D and 3D hand-pose estimation): distance between the thumb and pinky fingers tips.
- Hand movements (using 2D and 3D hand-pose estimation): averaged distance between each finger tip and the wrist point.
- Pronation-supination movements of the hands (using 3D hand-pose estimation): azimuthal angle from spherical coordinates of the tip of the thumb (computed with z-axis being the line between wrist and start of the third finger).

Each parameter was evaluated for all timesteps, smoothed using a Savitzky-Golay filter (with parameters window_length=9, polyorder=3) and then normalized between 0 and 1. Speed (in each direction) was also computed, smoothed using a Savitzky-Golay filter (with parameters window_length=5, polyorder=3) and normalized between −1 and 1.

The temporal evolution of the parameters was periodic because the protocols consisted of 10 repetitions of each movement. We used an algorithm based on *scipy*.*signal find_peaks* function to detect the 10 repetitions. Note that we restricted the analysis to the 9 repetitions that were clearly detectable and which did not depend on the initial position of the hand, since the aim was to study the whole dynamics of each sweep. The frequency *F* of the repetitions was measured. These 9 sweeps were analyzed individually and the following properties were computed: (i) duration of the sweep (noted *T*_*sweep*_ [*i*] for each sweep *i*); (ii) amplitude of the sweep (maximum minus minimum values) (noted *A*_*sweep*_); (iii) speed (amplitude divided by duration, noted *S*_*sweep*_). The three first/last sweeps properties were compared to study if the movement was slowed down or altered in any way during the protocol, i.e, we computed

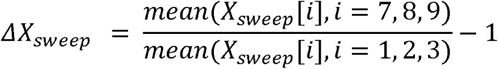

A fatigue parameter was also computed,

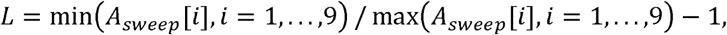

*L* represents the maximal change in amplitude during the whole protocol.

To compute the deviation from a periodic trajectory, we fitted the following function on each trace, using the *lmfit* package:

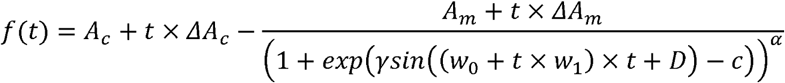

From these fitted parameters, we computed:

- The period variation, which represents the change in period computed from the fit parameters,

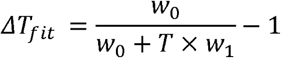

where *T* is the duration of the 9 sweeps.
- The amplitude variation, which represents the change in amplitude computed from the fit parameters,

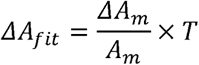

We used 7 parameters (*F, ΔT*_*sweep*_, *ΔA*_*sweep*_, *ΔS*_*sweep*_, *ΔT*_*fit*_, *ΔA*_*fit*_, *L*) for correlation with MDS-UPDRS III scores.

Videos were discarded for three reasons: (i) the algorithms DeepLabCut or HandGraphCNN failed to perform hand-pose estimation, the automated analysis failed (ii) to detect 9 sweeps or (iii) do the fitting procedure. Overall, 67% (64/96) of the videos for the tapping finger, 83% (78/94) for the hand movements, 43% (35/82) for the pronation-supination movements were used for further analysis.

### Impact of the measured parameters for MDS-UPDRS scoring using statistical learning algorithms

To evaluate the impact of the measured parameters for MDS-UPDRS scoring, three classical algorithms of machine learning were tested: (i) the linear regression, (ii) the decision tree with max_depth=2 and (iii) the decision tree with max_depth=3, all from the *scikit-learn* toolbox. We trained these algorithms on the task of predicting the averaged MDS-UPDRS III score, from the 7 previously defined parameters, for each protocol.

The coefficient of determination after training was computed with correct labels. Concurrently, we estimated the distribution of coefficients of determination obtained when training the algorithm on shuffled MDS-UPDRS III scores: we computed the mean and standard deviation on 100 different shuffles, and approximated them with a normal distribution. Using this control distribution, we estimated the probability *p*_*shuffle*_ that trained networks with shuffled scores had higher coefficients of determination than the one obtained with the correct MDS-UPDRS scores. We used *p*_*shuffle*_ to study the significativity of the predictions (*: *p*_*shuffle*_ <0.05, **: *p*_*shuffle*_ <0.005, ***: *p*_*shuffle*_ <0.0005).

Finally, individual correlations were computed between the averaged MDS-UPDRS III scores and the 7 different parameters computed in the previous section, using *scipy*.*stats linregress* and *spearmanr* functions. Slopes and p-values are extracted for each case. Results with p < 0.05 were considered statistically significant (*: p<0.05, **: p<0.005, ***: p<0.0005).

## RESULTS

### 1. Demographic features

Thirty-six parkinsonian patients were enrolled from the Movement Disorder Unit at Avicenne University Hospital, between June 2019 and December 2020 (Table S1). Thirty-four patients had a confirmed PD diagnosis according to MDS clinical criteria, 2 of them having a genetic form[1]. Among the 2 remaining patients, one fulfilled the criteria of Multi System Atrophy and the other one had a progressive atypical parkinsonism of undetermined origin. Patients’ assessment was carried out without considering the last dopamine intake. We also recruited 11 healthy individuals, without any known neurological condition, to assess bradykinesia in *priori* non-parkinsonian subjects.

### 2. Unreliability of MDS-UPDRS III scoring for bradykinesia

Five neurologists were asked to score N=272 videos from PD and non-PD subjects performing the 3.4, 3.5 and 3.6 items of the MDS-UPDRS III, with both hands. Comparing the different scores and analyzing their distribution for each video led to the conclusion that in the majority cases, the five examiners did not give the same scores. We computed the standard deviation of the scores for each video and plotted its distribution (Figure S1a). We observed that 58 videos had a null standard deviation, and as such were scored similarly by all five neurologists. When considering the whole dataset, MDS-UPRDS scores had a mean standard deviation of 0.409.

We also gathered videos as a function of the number of different scores that were given by the neurologists, and then in each class on the difference between the maximum and minimum scores (Figure S1b). 58 videos had the same ratings, 171 with 2 different scores, 42 with 3 different scores and even 1 with 4 different scores given for the same videos (on a total of 5 possible scores). Overall, this confirms the existence of inter-rater variability for MDS-UPDRS III scoring.

### 3. Extractions of relevant parameters for MDS-UPDRS III scores using deep learning

Using deep learning algorithms (DeepLabCut and HandGraphCNN), we detected 21 important points describing the hands of the patients, and extrapolated their coordinates in 2D and 3D as a function of time (Methods).

After complex steps of post-processing and analysis, one metric is extracted for each protocol and represented across time. Examples of extracted data in a single patient are presented in Figures 1, 2 and 3 for finger tapping, hand movements and pronation-supination movements of the hands, respectively.

**Figure 1.**
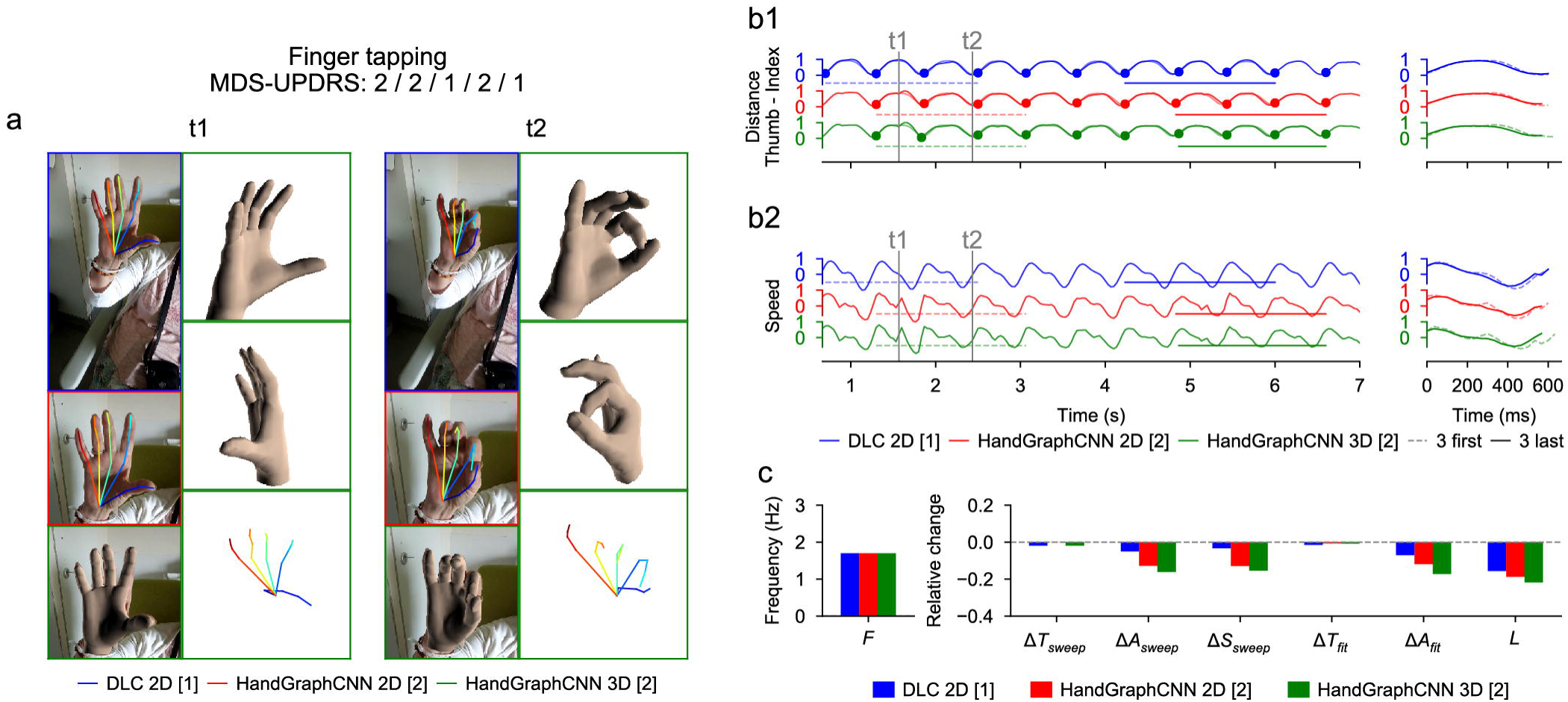
Video analysis using deep learning algorithms for tapping protocol (associated Supplementary Movie 1). (a) Snapshots from the initial videos, with the different skeletons extracted using the two deep learning algorithms (blue DeepLabCut 2D, red HandGraphCNN 2D, green HandGraphCNN 3D). (b1, left) Evolution of the distance between the thumb and pinky fingers tips with 9 sweeps (delimited by the 10 plain circles on each trajectory). (b1, right) First three sweeps (dotted lines) compared to the last three ones (plain lines). (b2) same as (b1) for the speed. (c) Frequency *F*, fatigue *L*, relative variation of the period Δ*T*_*sweep*_, amplitude Δ*A*_*sweep*_, speed Δ*S* _*sweep*_ accross the protocol, more complex variation coefficients Δ*T*_*fit*_ and Δ*A*_*fit*_ based on the fit of a periodic function.

**Figure 2.**
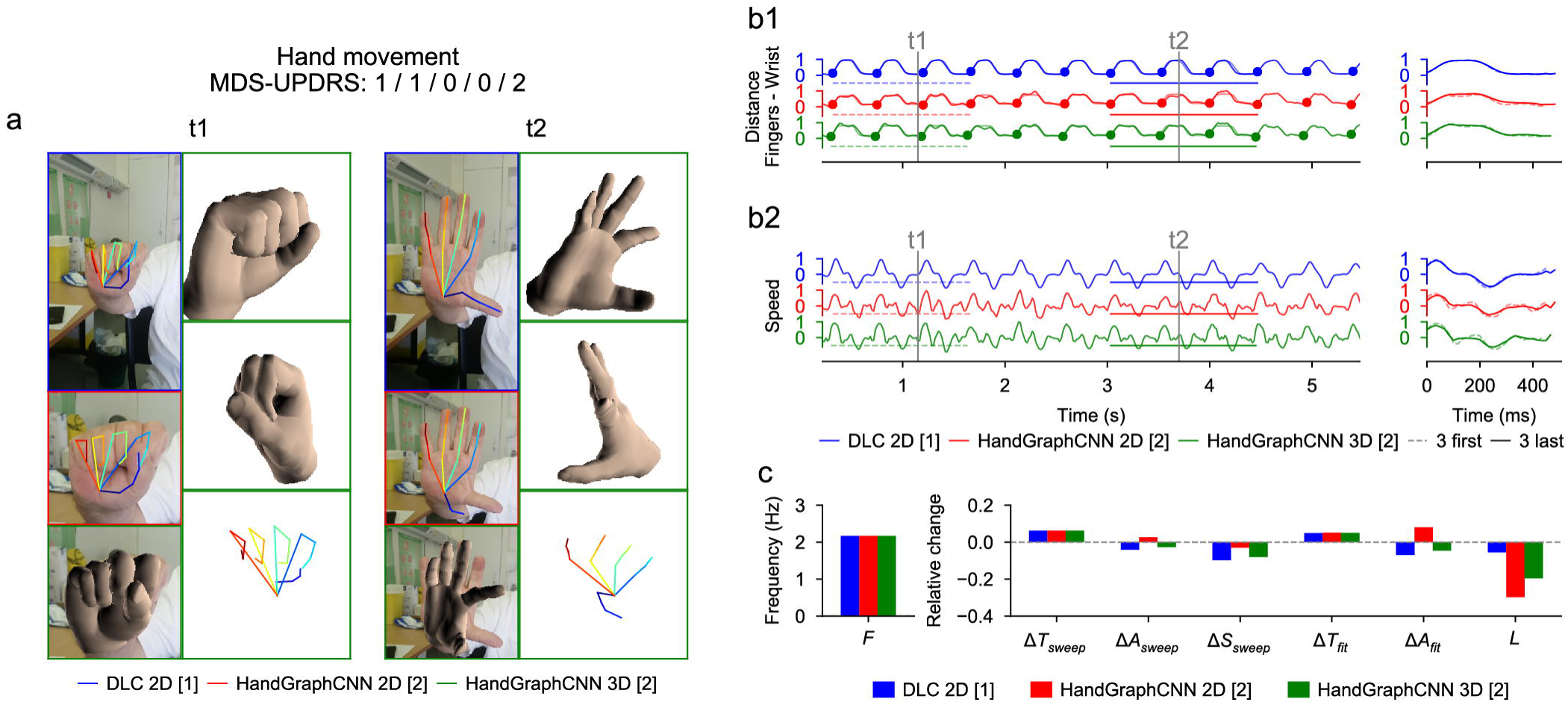
Video analysis using deep learning algorithms for hand movements protocol (associated Supplementary Movie 2). (a) Snapshots from the initial videos, with the different skeletons extracted using the two deep learning algorithms (blue DeepLabCut 2D, red HandGraphCNN 2D, green HandGraphCNN 3D). (b1, left) Evolution of the averaged distance between each finger tip and the wrist point with 9 sweeps (delimited by the 10 plain circles on each trajectory). (b1, right) First three sweeps (dotted lines) compared to the last three ones (plain lines). (b2) same as (b1) for the speed. (c) Frequency *F*, fatigue *L*, relative variation of the period Δ*T*_*sweep*_, amplitude Δ*A*_*sweep*_, speed Δ*S* _*sweep*_ accross the protocol, more complex variation coefficients, Δ*T*_*fit*_ and Δ*A*_*fit*_ based on the fit of a periodic function.

**Figure 3.**
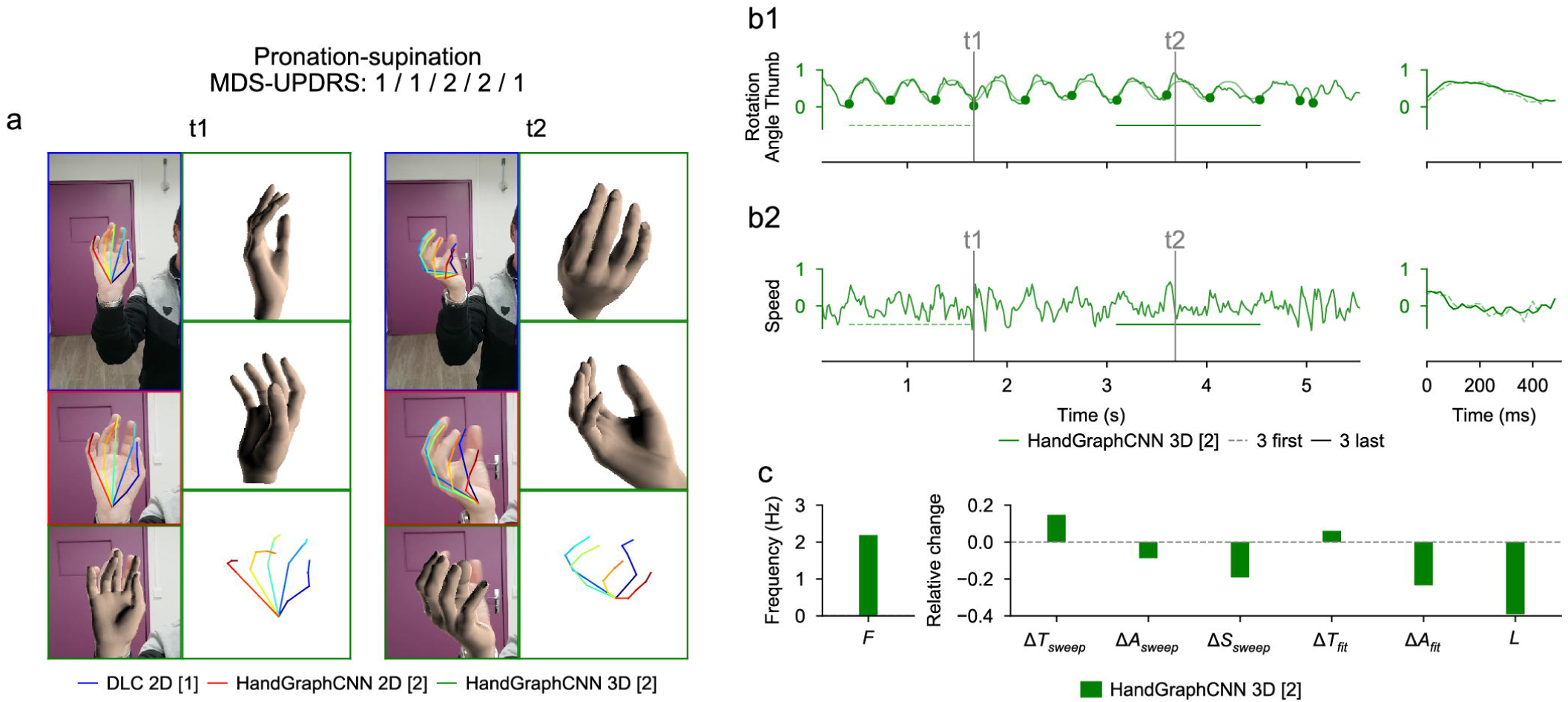
Video analysis using deep learning algorithms for prono-supination protocol (associated Supplementary Movie 3). (a) Snapshots from the initial videos, with the different skeletons extracted using the two deep learning algorithms (blue DeepLabCut 2D, red HandGraphCNN 2D, green HandGraphCNN 3D). (b1, left) Evolution of the azimuthal angle from spherical coordinates of the tip of the thumb with 9 sweeps (delimited by the 10 plain circles on each trajectory). (b1, right) First three sweeps (dotted lines) compared to the last three ones (plain lines). (b2) same as (b1) for the speed. (c) Frequency *F*, fatigue *L*, relative variation of the period Δ*T*_*sweep*_, amplitude Δ*A*_*sweep*_, speed Δ*S* _*sweep*_ accross the protocol, more complex variation coefficients, Δ*T*_*fit*_ and Δ*A*_*fit*_ based on the fit of a periodic function.

For each Figure, we present snapshots from the initial videos, with the different skeletons extracted using the two deep learning algorithms (Figures 1a/2a/3a, blue DeepLabCut 2D, red HandGraphCNN 2D, green HandGraphCNN 3D). From the trajectories of the hand joints, we computed protocol specific metrics (Methods) during the whole protocol. As the protocol is composed of 10 repetitions of the same hand movement, for each metric 9 sweeps (delimited by the 10 plain circles on each trajectory, see Figures 1b1/2b1/3b1, left) were detected and then analyzed with the evolution of movement during the protocols. The shapes from the first three sweeps (dotted lines) were compared to the last three ones (plain lines) (Figures 1b1/2b1/3b1, right). Similarly, we computed the associated speed in Figures 1b2/2b2/3b2. The frequency *F*, fatigue *L*, relative variation of the period Δ*T*_*sweep*_, amplitude Δ*A*_*sweep*_ and speed Δ*S*_*sweep*_ accross the protocol were computed and presented in Figures 1c/2c/3c. More complex variation coefficients, Δ*T*_*fit*_ and Δ*A*_*fit*_ based on the fit of a periodic function were also obtained (Methods) and represented in Figures 1c/2c/3c.

For the finger tapping protocol (Figure 1), we computed the distance between the tips of the thumb and the index fingers as the primary metric, and the associated speed. We observed on this example that the trajectories were almost periodic. The analysis showed that, with the three different detection algorithms (Figure 1: DeepLabCut 2D (blue), HandGraphCNN 2D (red) and HandGraphCNN 3D (green)), the parkinsonian patient reduced the amplitude Δ*A*_*sweep*_/Δ*A*_*fit*_ of its movement during the protocol, and also reduced its speed. Δ*S*_*sweep*_ The period Δ*T*_*sweep*_ /Δ*T*_*fit*_ stayed constant during the experiment. This agrees with the slightly positive MDS-UPDRS III scores (2/2/1/2/1) given by the neurologists.

For the hand movements (Figure 2), we computed the mean distance between the finger tips and the wrist. We observed a periodic trajectory, with an increase of the period Δ*T*_*sweep*_/Δ*T*_*fit*_, and consequently a reduction of the speed Δ*S* _*sweep*_. Since period is harder to assess than amplitude, it might explain why the raters gave different scores (1/1/0/0/2).

For the pronation-supination movements of the hands (Figure 3), an angle was computed from the 3D representation. The trajectories were periodic enough to enable detection of the sweeps. Because of the irregularity of the trajectory, it is more accurate to use measures from fits to a periodic function. There was a drop of the amplitude of the movement Δ*A*_*fit*_, consistent with the MDS-UPDRS III scores (1/1/2/2/1).

Overall, these three examples highlight the potential of the current analysis to quantify hand movements during MDS-UPDRS III protocols.

### 4. Impact of the measured parameters on MDS-UPDRS using statistical analysis

We trained three algorithms (linear regression, decision tree with max_depth=2, decision tree with max_depth=3) to predict the averaged MDS-UPDRS III score from the previously defined 7 variables for each video. The results are presented in Table 1 with the coefficients of determination obtained for algorithms trained with correct MDS-UPDRS scores, the averaged coefficients of determination over 100 different randomly shuffled datasets and the *p*_*shuffle*_ value used for significativity (see Methods for its definition). For the finger tapping and hand movements protocols, all 3 algorithms predicted significantly better the correct MDS-UPDRS III score than algorithms trained with shuffled scores (Table 1). For the prono-supination movements of the hands protocol, algorithms failed to predict the correct scores since algorithms trained with the correct scores performed as well as the ones trained with shuffled scores (*p*_*shuffle*_ >0.1). Using more complex algorithms (Linear regression < Decision tree with max_depth=2 < Decision tree with max_depth=3) led to higher coefficient of determinations for each protocol, but to decreases in significativity (Table 1).

**Table 1.**
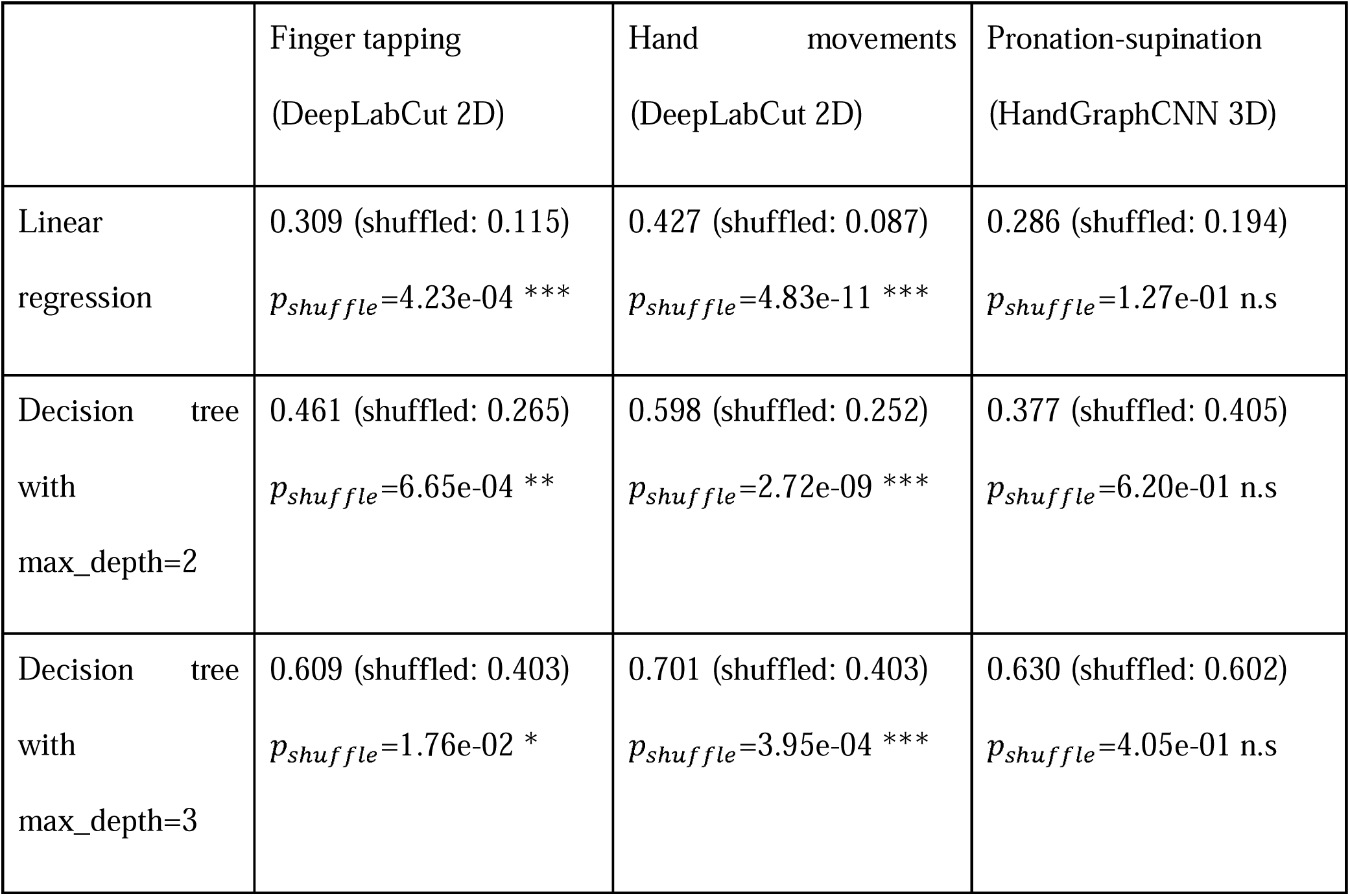
Regression algorithms trained to match MDS-UPDRS scores based on the parameters detected by the automated analysis. Coefficients of determination are presented for training with the correct scores, the averaged over 100 trainings with shuffled scores, and the probability *p*_*shuffle*_ used to test significativity (see Methods).

|  | Finger tapping<br>(DeepLabCut 2D) | Hand movements<br>(DeepLabCut 2D) | Pronation-supination<br>(HandGraphCNN 3D) |
| --- | --- | --- | --- |
| Linear regression | 0.309 (shuffled: 0.115)<br>$p_{shuffle}=4.23e-04$ *** | 0.427 (shuffled: 0.087)<br>$p_{shuffle}=4.83e-11$ *** | 0.286 (shuffled: 0.194)<br>$p_{shuffle}=1.27e-01$ n.s |
| Decision tree<br>with<br>max_depth=2 | 0.461 (shuffled: 0.265)<br>$p_{shuffle}=6.65e-04$ ** | 0.598 (shuffled: 0.252)<br>$p_{shuffle}=2.72e-09$ *** | 0.377 (shuffled: 0.405)<br>$p_{shuffle}=6.20e-01$ n.s |
| Decision tree<br>with<br>max_depth=3 | 0.609 (shuffled: 0.403)<br>$p_{shuffle}=1.76e-02$ * | 0.701 (shuffled: 0.403)<br>$p_{shuffle}=3.95e-04$ *** | 0.630 (shuffled: 0.602)<br>$p_{shuffle}=4.05e-01$ n.s |

In conclusion, the different parameters computed using our automated analysis include pertinent information for 3.4 and 3.5 MDS-UPDRS scoring.

### 5. Individual correlations with MDS-UPDRS III

Figure 4 showed correlations of the measured parameters with the averaged MDS-UPDRS III scores. For the finger tapping protocol (top), metrics extracted from the 2D DeepLabCut coordinates are shown here and their correlation with MDS-UPDRS III scores is presented. The variation of amplitude (for both empirical measure and fit), speed and fatigue are significantly correlated with the MDS-UPDRS III scores, with a negative slope, as expected from MDS-UPDRS III guidelines. The other metrics (frequency and period variations) are not significantly correlated. Similar results are observed for the hand movements protocol (center), with higher significant correlations for amplitude (both types of measure), speed and fatigue. Moreover, the period variation measured with the fit is also negatively and significantly correlated. This analysis did not reach significance for pronation-supination movements of the hands (bottom). Overall, measured parameters related to MDS-UPDRS guidelines are directly correlated with MDS-UPDRS scores.

**Figure 4.**
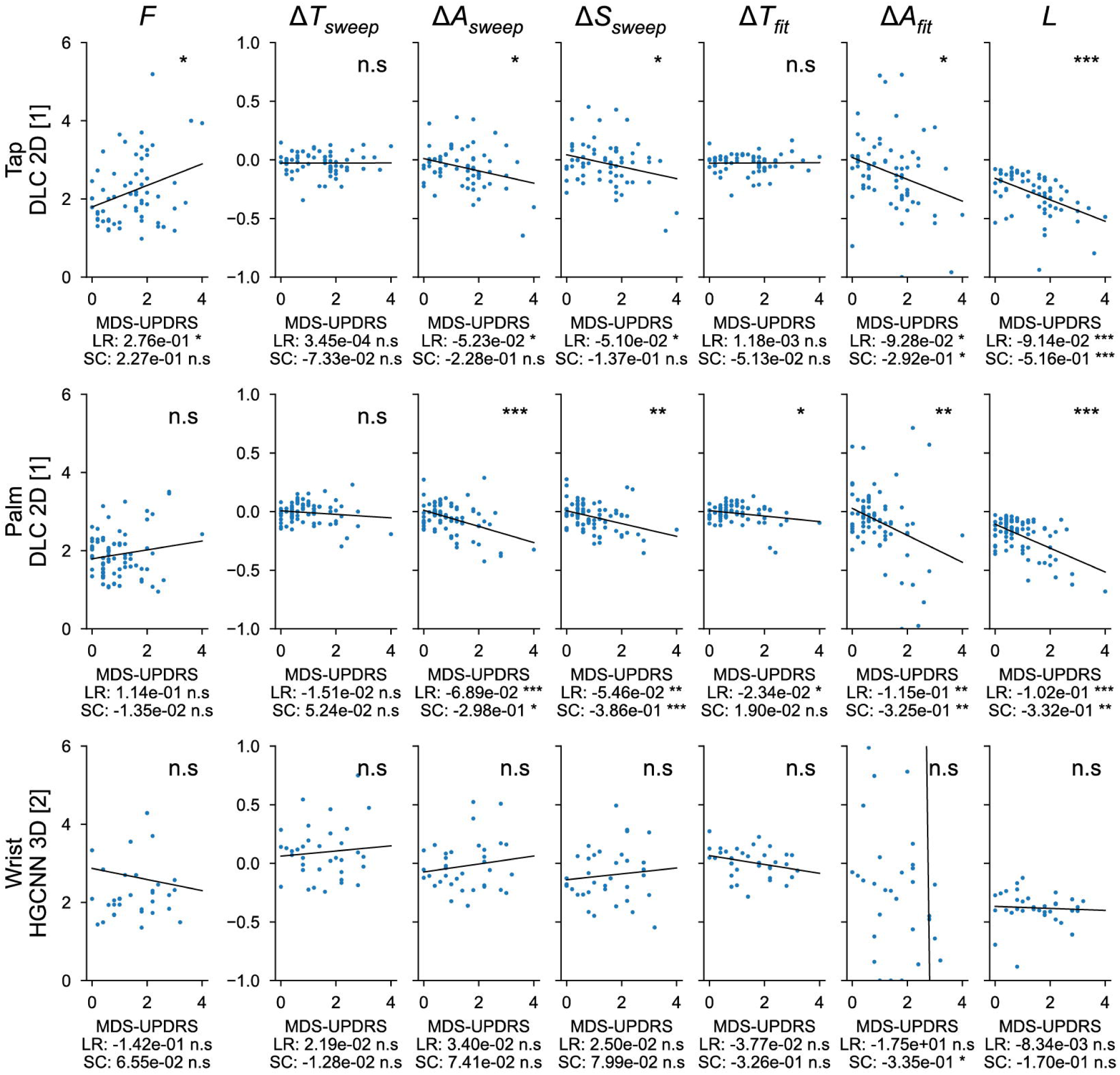
Correlations between the 7 metrics computed with deep learning algorithms and averaged MDS-UPDRS scores. *F*: frequency of the repetitions; Δ*T*_*sweep*_: comparison of the duration of the three first/last sweeps; Δ*A*_*sweep*_: comparison of the amplitude of the three first/last sweeps; Δ*S*_*sweep*_: comparison of the speed of the three first/last sweeps; Δ*T*_*fit*_: period variation, which represents the change in period computed from the fit parameters; Δ*A*_*fit*_: amplitude variation, which represents the change in amplitude computed from the fit parameters; *L*: fatigue parameter which represents the maximal change in amplitude during the whole protocol. ns: not significant;* <0.05, **<0.005, ***<0.0005 for the significance of the p-values.LR: linear regression; SC: Spearman correlation.

## DISCUSSION

In this study, we created an innovative tool able to reproduce clinical observations obtained with MDS-UPDRS scores with a precise and objective quantification and an adequate evaluation of the MDS-UPDRS III scores for finger tapping and hand movements. We obtained an accurate extraction of important temporal characteristics of MDS-UPDRS III hand tasks, in an automated way only starting from a standard video.

The 7 parameters tested here, exploring speed, frequency, amplitude, duration and fatigue were measured using the automated analysis for finger tapping and hand movements protocols, predicted the MDS-UPDRS III used worldwide for PD motor symptoms evaluation with high coefficients of determination. More importantly, predictions using different statistical learning regression algorithms were significantly higher than algorithms trained with shuffled datasets. We concluded that the 7 parameters computed in the analysis presented here contained enough information for MDS-UPRDS estimation for finger tapping and hand movement protocols. For pronation-supination movements of the hands, our methods did not lead to statistically significant predictions. This might be explained by the low number of samples to reach a significant response. Indeed, less than half of the videos with prono-supination movements of the hands passed all the tests to be included in the prediction analysis. Moreover, we computed individual correlations for the three protocols, and showed that the variation of amplitudes, speed and fatigue were significantly correlated with the MDS-UPRDS III scores. Importantly, the parameters measured in this analysis were consistent with MDS-UPRDS III guidelines for scoring. The 2D and 3D models of the hand recorded by video allow precise quantification of multiple parameters such as speed, amplitude and rate of the movement during MDS-UPDRS III evaluation. Thus, the composite parameters analyzed here are the same as those used for assessing bradykinesia during a medical consultation. Bradykinesia is a complex phenomenon of slowness of movement that cannot be seen only as a simple decrease of the movement rate [20], and therefore more complex properties need to be considered in the scoring. The video-based method described in our study by-passes the subjective measurements of these parameters.

As already stated in the literature [7–12], we also highlight here that MDS-UPDRS III scoring is physician-dependent, and as such is a less reliable parameter than an automated and quantifiable assessment tool to evaluate bradykinesia. Here, we video-taped patients in real-life situations without specific preparation and environment. Moreover, patients were not selected regarding the phenotype or the severity of the parkinsonian syndrome. Importantly, by labelling only 5 frames per videos we demonstrate that, even with scarce labels, the network performs accurately. Overall, the time needed to label 5 frames is quite small (1-2 minutes for an expert) while the operation would last at least an hour to individually label each frame, leading to a great gain in term of time for a result almost as precise as manual labelling.

In the expanding field of telemedicine, the present tool appears of much interest. As an example, COVID-19 pandemic has been striking evidence that remote evaluation of neurological patients is needed, this being even more crucial with chronic diseases [27–29]. It has been shown that remote assessment of MDS-UPDRS III is feasible via videoconferencing, except for specific items such as rigidity, or postural instability, which are not as important as bradykinesia for general evaluation of the disease [30,31]. Indeed, some studies, which restricted bradykinesia assessment to the upper limb motor tasks of the MDS-UPDRS III, showed that upper limb motor performance was a predictive feature of PD onset and progression[22,32]. Research on remote evaluation of PD patients has been mainly focused on the use of technological devices [14,15,33,34]. Such techniques require specific setup, which differs from MDS-UPDRS III tasks (e.g., touching tactile screen of tapping, holding the phone during prono-supination). Recording video while clinically assessing the MDS-UPDRS III is easy, and therefore any videos recorded while following MDS-UPDRS III instructions can be analyzed and quantified by our system. Also, it is well known that PD patients’ symptoms, such as tremor and motor functions, vary upon emotional load [35,36]. Thus, by quantifying bradykinesia from homemade videos, in a less stressful environment than hospital or medical consultation, clinicians could have a more reliable assessment of their patient’s condition on a daily life basis. It is noteworthy that this deep-learning analysis of parkinsonian movement could be extended to other body parts (e.g., feet movements, hypomimia, posture) and therefore most of the MDS-UPDRS III procedures. Thus, a wide range of movement disorders, such as tremor or chorea, could be eligible to this strategy of evaluation [16,37].

This study has several limitations. First, it is a monocentric study, which needs to be extended to other centers. Nevertheless, our PD patient population was similar to epidemiological data found in the medical literature. Secondly, there was an inter-rater variability in the MDS-UPDRS part III scoring which cannot be predicted by the deep-learning analysis. To circumvent this limitation, we performed five assessments by different raters, blinded to each other, for each test on every PD patients and control subjects. This allowed us to reduce this variability and limit measurement errors. Thirdly, regarding pronation-supination movements of the hands, the results were not significant despite a trend for the amplitude parameter. This could be explained by the low number of patients but also the complexity of extracting 3D coordinates. Frontal video recording of the hand pronation-supination movements with the forearm horizontal and not vertical would certainly facilitate the hand rotation analysis. Further prospective analyses with more patients are needed to implement this specific assessment. Forthcoming development of this software will allow analysis of other movement disorders and other body parts.

In conclusion, using a deep-learning approach, we provided a quantitative measurement of bradykinesia that prevents from inter-operator variability. We have reached an unprecedented level of precision and simplicity for its assessment. This precision, even at distance, could help non-movement disorder specialists to rate bradykinesia of their patients accurately. It would also be useful for specialists and non-specialists, to monitor bradykinesia of patients at distance, with video recordings provided directly by the patient or caregivers, with appropriate instructions.

## Acknowledgment

We would thank all the patients for accepting to be filmed.

## Author’s Roles

1. Research project: A. Conception, B. Organization, C. Execution
2. Statistical analysis: A. Design, B. Execution, C. Review and critique
3. Manuscript: A. Writing of the first draft, B. Review and critique.

GV : 1A, 1B, 1C, 2A, 2B, 3A, 3B

CD : 1B, 1C, 3A, 3B

QS : 1B, 1C, 3A, 3B

MM : 1C, 3B

BG : 1C, 3B

LV : 1A, 2C, 3B

BD : 1A, 1B, 1C, 2C, 3B

## Financial Disclosure/Conflict of interest

GV, CD, QS, MM, BG, LV declare no conflicts of interest. BD received research support from Orkyn, Merz-Pharma and Contrat de Recherche Clinique 2021 (CRC 2021). No sponsorship was obtained for this study.

## Supplementary Figure

**Figure S1. Unreliability of MDS-UPDRS scores across neurologists’ ratings.** (**a**) Histogram of the standard deviations (computed for each video, on the 5 different MDS-UPRS scores); (**b**) videos ranked by the number of different MDS-UPDRS scores given by the physicians, colors for the difference between maximum and minimum scores.

## Notes

### Competing Interest Statement

The authors have declared no competing interest.

