## Supplementary figures and images for "Video-based automated analysis of MDS-UPDRS III parameters in Parkinson disease"

### Supplemental Figure 1

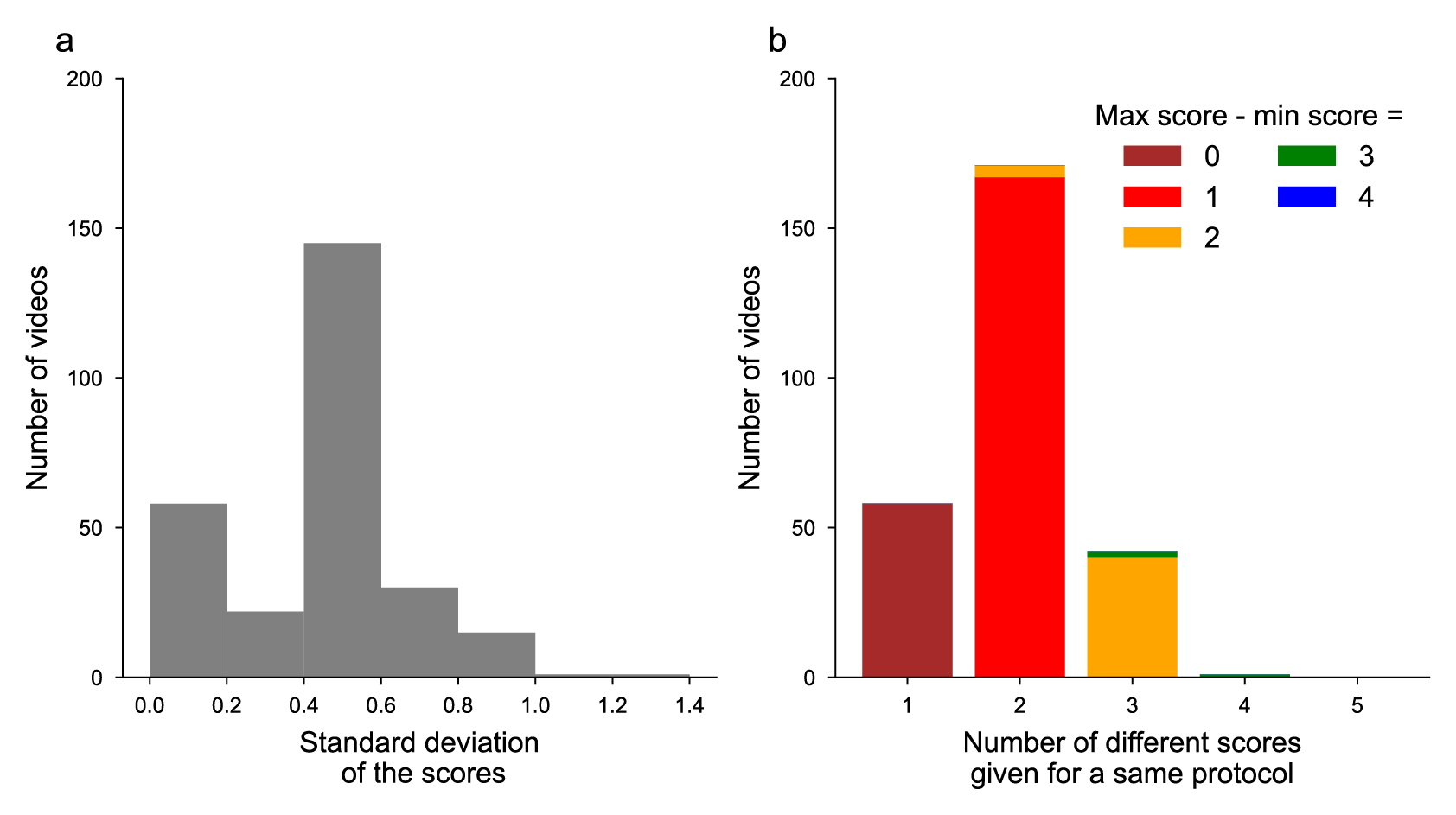
