## Supplemental Table 1 for "Video-based automated analysis of MDS-UPDRS III parameters in Parkinson disease"

**Table S1. Patients’ clinical and demographic characteristics.**

| **Gender (%)** |  |
| --- | --- |
| Male | 28 (77.8%) |
| Female | 8 (22.2%) |
| **Age (mean; min-max)** | 61.2 (32.7-80.1) |
| **Disease duration in months (mean; min-max)** | 95.9 (269.7-3.5) |
| **Parkinson’s Disease (%)** | 32/36 (88.9%) |
| **Others parkinsonian syndromes** |  |
| Atypical | 2/36 (5.5%) |
| Genetic | 2/36 (5.5%) |
| **Most affected side (%)** |  |
| Left | 14/36 (38.9%) |
| Right | 21/36 (58.3%) |
| Axial | 1/36 (3.8%) |
| **Amount of dopamine in mg (mean; min-max)** | 560.6 (0-1317) |
| **Deep Brain Stimulation (%)** | 5/36 (13.9%) |
| **MDS-UPDRS score part III (median; min-max)** | 6 (0-28) |
| **Hoehn & Yahr score (median; min-max)** | 2 (0-3) |
